# Deuterated rhodamines for protein labelling in nanoscopy

**DOI:** 10.1101/2020.08.17.253880

**Authors:** Kilian Roßmann, Kerem C. Akkaya, Corentin Charbonnier, Jenny Eichhorst, Ben Jones, Martin Lehmann, Johannes Broichhagen

## Abstract

Rhodamine molecules are setting benchmarks in fluorescence microscopy. Herein, we report the deuterium (d12) congeners of tetramethyl(silicon)rhodamine, obtained by isotopic labelling of the four methyl groups, which improves photophysical (*i.e.* brightness, lifetimes) and chemical (*i.e.* bleaching) properties. We explore this finding for SNAP- and Halo-tag labelling, and highlight enhanced properties in several applications, such as Förster resonance energy transfer, fluorescence activated cell sorting, fluorescence lifetime microscopy and stimulated emission depletion nanoscopy. We envision deuteration as a generalizable concept to improve existing and develop new Chemical Biology Probes.

## MAIN TEXT

Fluorescence microscopy is the technique of choice in modern biomedical research to elucidate structures or to interrogate function. Efforts to improve performances resulted in the development of super-resolution microscopy (nanoscopy), which is defined to obtain resolution higher than the diffraction limit described by Abbe’s law.^[1]^ While optics and instruments have been advanced constantly, the elaboration of synthetic molecular dyes has driven the field to the current state-of-art.^[2]^ Given their small size, and the possibility to target them to cellular organelles (by means of substitution patterns) or proteins (by means of self-labelling tags, *e.g.* SNAP/Halo-tag fusions) make them an attractive choice for the visualization and interrogation of biomolecular function.^[3]^ Several chemical scaffolds, for instance nitrobenzodioxazoles (NBDs)^[4]^ and coumarins^[5]^, remain interesting synthetic targets, yet xanthene dyes, which include tetramethylrhodamine (TMR) and silicon rhodamine (SiR), experienced a renaissance in terms of novel modifications to tune and boost important parameters, such as color, brightness, lifetime and reactivity.^[6–10]^ However, and best to our knowledge, isotopic labelling has not been among this repertoire.

We fill this gap by designing, synthesizing, and testing tetramethylrhodamines, in which each CH_3_ group is replaced by CD_3_, and furthermore equipped with a bioorthogonal handle for SNAP- or Halo-tag labelling of fused proteins (Fig. 1A, Scheme S1). Comparing deuterated and conventional rhodamines, we intuitively find that some photophysical properties remained unchanged, such as maximal excitation and emission wavelengths (λ_Ex/Em_ (TMR(-d12)) ∼ 549/576 nm; λ_Ex/Em_ (SiR(-d12)) ∼ 650/670 nm) (Fig. 1A, B), while others do change dramatically, such as extinction coefficients (ε(TMR) = 78,000^[6]^ *vs.* ε(TMR-d12) = 83,000 M^−1^ cm^−1^; ε(SiR) = 141,000^[6]^ *vs.* ε(SiR-d12) = 139,000 M^−1^ cm^−1^) and quantum yields (Φ(TMR) = 0.43 *vs.* Φ(TMR-d12) = 0.55; Φ(SiR) = 0.41^[11]^ *vs.* SiR-d12 = 0.57), leading to augmented brightness (ε × Φ: TMR = 34 *vs.* TMR-d12 = 46; SiR = 0.58 *vs.* SiR-d12 = 0.79) (Fig. 1A). Chemical fusion to *O*^6^-benzylguanine or a chloroalkane linker yields labelling substrates for the SNAP- and Halo-tag, respectively. The *in vitro* labelling of SNAP with BG-TMR-d12 and BG-SiR-d12 could be traced by an increase in fluorescence polarization to determine labelling kinetics, which do not differ between hydrogenated and deuterated substrates (t_1/2_(TMR) = 13.3 *vs.* t_1/2_(TMR-d12) = 12.2 s; t_1/2_(SiR) = 23.5 *vs.* t_1/2_(SiR-d12) = 29.4 s) (Fig. 1A, C). By labelling a SNAP-Halo construct with Halo-TMR/BG-SiR and BG-TMR/Halo-SiR, the improved photophysical properties lead to an increased efficiency in Förster resonance energy transfer (FRET) by 2% and 8%, respectively, for the d12 variants (Fig. 1D). These results highlight that even subtle chemical changes can have pronounced effects on spectroscopic properties, exploring the chemical space in a new direction.

**Figure 1.**
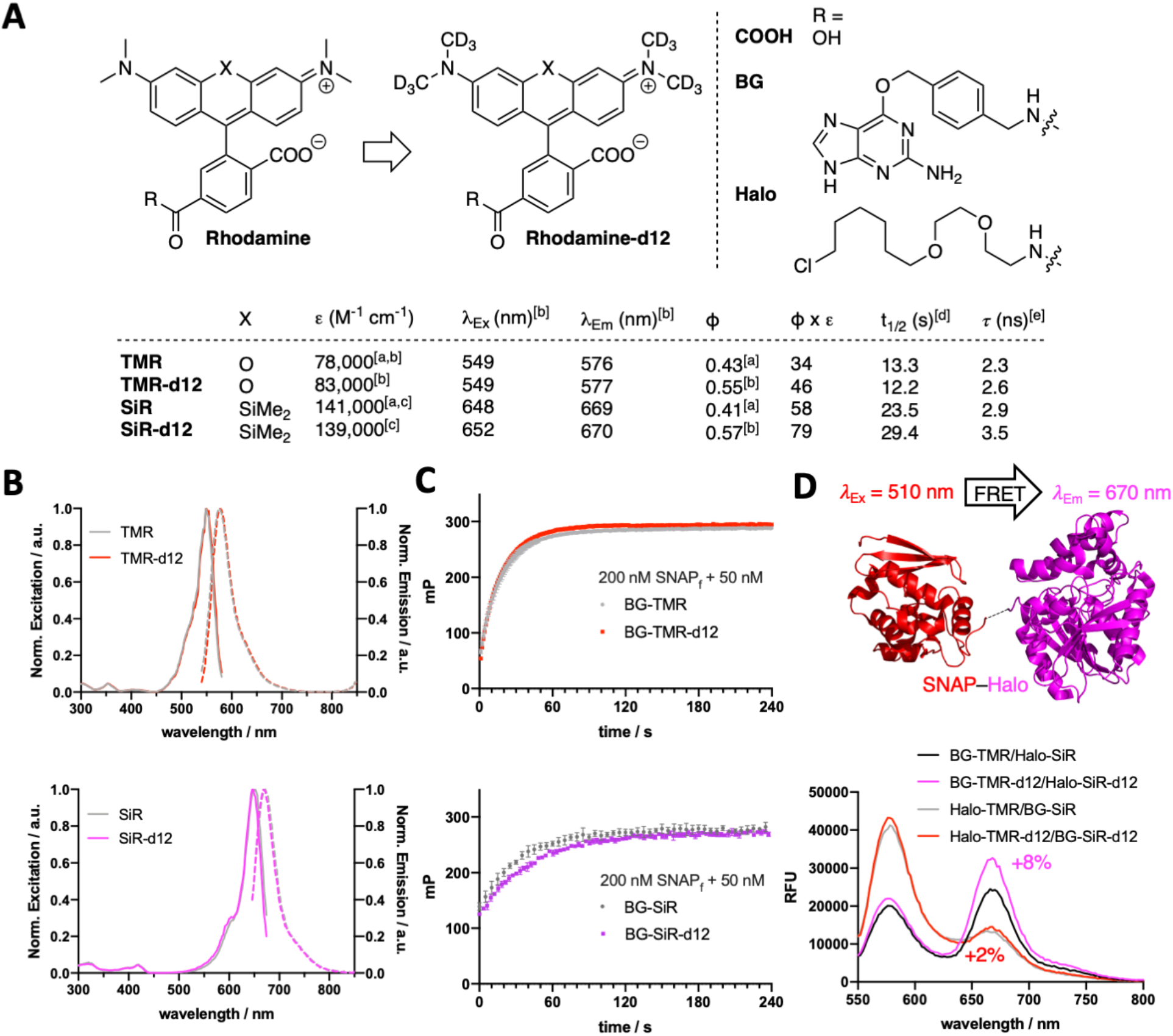
Deuteration improves rhodamine brightness. A) The isotopic H/D exchange on the four methyl groups leads to d12 variants of TMR (X = O) and SiR (X = SiMe_2_). Derivatization on the 6-carboxylate allows synthesis of BG- and Halo-congeners for SNAP- and Halo-tag labelling, respectively. B) Excitation and emission spectra of TMR(-d12) (top) and SiR(-d12) (bottom). C) Fluorescence polarization (mP) assay of BG-TMR(-d12) (top) and BG-SiR(-d12) (bottom) when incubated with SNAP_f_ to determine labelling kinetics. D) *In vitro* FRET assay of a BG-TMR(-d12) and Halo-SiR(-d12) labelled SNAP-Halo protein shows an increase of acceptor/donor emission ratio. ^[a]^ reference values; ^[b]^ in activity buffer; ^[c]^ in EtOH + 1% TFA; ^[d]^ R = BG and SNAP_f_ *in vitro* in activity buffer; ^[e]^ SNAP bound in cells.

We next turned to cellular labelling and imaging and aimed to determine the behavior of our dyes on targets that are mainly expressed on the membrane, as well as targets that are found intracellularly, both regions that have been addressed for SNAP-tag labelling.^[3]^ First, we employed CHO-K1 cells stably expressing SNAP-tagged glucagon-like peptide 1 receptor (SNAP-GLP1R:CHO-K1)^[12]^, a cell line intensely used to study the physiology of this blockbuster class B G protein-coupled receptor (GPCR) involved in glucose homeostasis and targeted in diabetes treatment^[13,14]^, as a benchmark for d12 performances. As such, cells were labelled with 1 μM BG-TMR/SiR(-d12) for 30 min, before washing and live imaging by confocal microscopy, which revealed staining of SNAP-GLP1R with all deuterated and parental dyes tested (Fig. 2A). Having established labelling on the outer plasma membrane, we secondly investigated intracellular staining in HeLa cells that stably express SNAP-tagged Cox8A (SNAP-Cox8A:HeLa), located in the inner mitochondrial membrane (Fig. 2B), which has been used to study mitochondrial ultrastructure in live cells.^[15,16]^ As for SNAP-GLP1R, we observed clean labelling with all dyes, and for both colors with an observable increase in brightness for the d12 derivatives. With this enhanced performance in microscopy, we wanted to quantify brightness by fluorescence activated cell sorting (FACS) to obtain robust values over large sample sizes. Accordingly, we labelled SNAP-GLP1R:CHO-K1 and SNAP-Cox8A:HeLa cells with both, BG-TMR(-d12) and BG-SiR(-d12) to compare red and far-red color intensities by subsequent sorting (Fig. 2C). Histograms of SNAP-GLP1R:CHO-K1 cells labelled showed a right-shift in fluorescence intensity when dyes were deuterated (Fig. 2C, left). In contrast, SNAP-Cox8A:HeLa cells were labelled more homogeneously with a pronounced shift to higher intensities for SiR-d12 compared to its non-deuterated congener (Fig. 2C, right), while TMR(d12) only displayed a subtle change. By normalizing intensities and comparison, we calculate higher mean intensities for our deuterated dye versions (Fig. 2D). While no large increase was observed in SNAP-Cox8A:HeLa cells for TMR-d12 (2%), mean intensity was markedly increased in SNAP-GLP1R:CHO-K1 cells (28%). Furthermore, SiR-d12 outperformed SiR on SNAP-GLP1R and SNAP-Cox8A with an intensity increase of 43% and 50%, respectively, which compares well with in vitro increase in brightness (35% and XX%). These results demonstrate that rhodamines with CD_3_ groups on the amines are not only the applicable to live cells but even outperform non-deuterated fluorophores, which is in line with our *in vitro* data of the unbound dyes.

**Figure 2.**
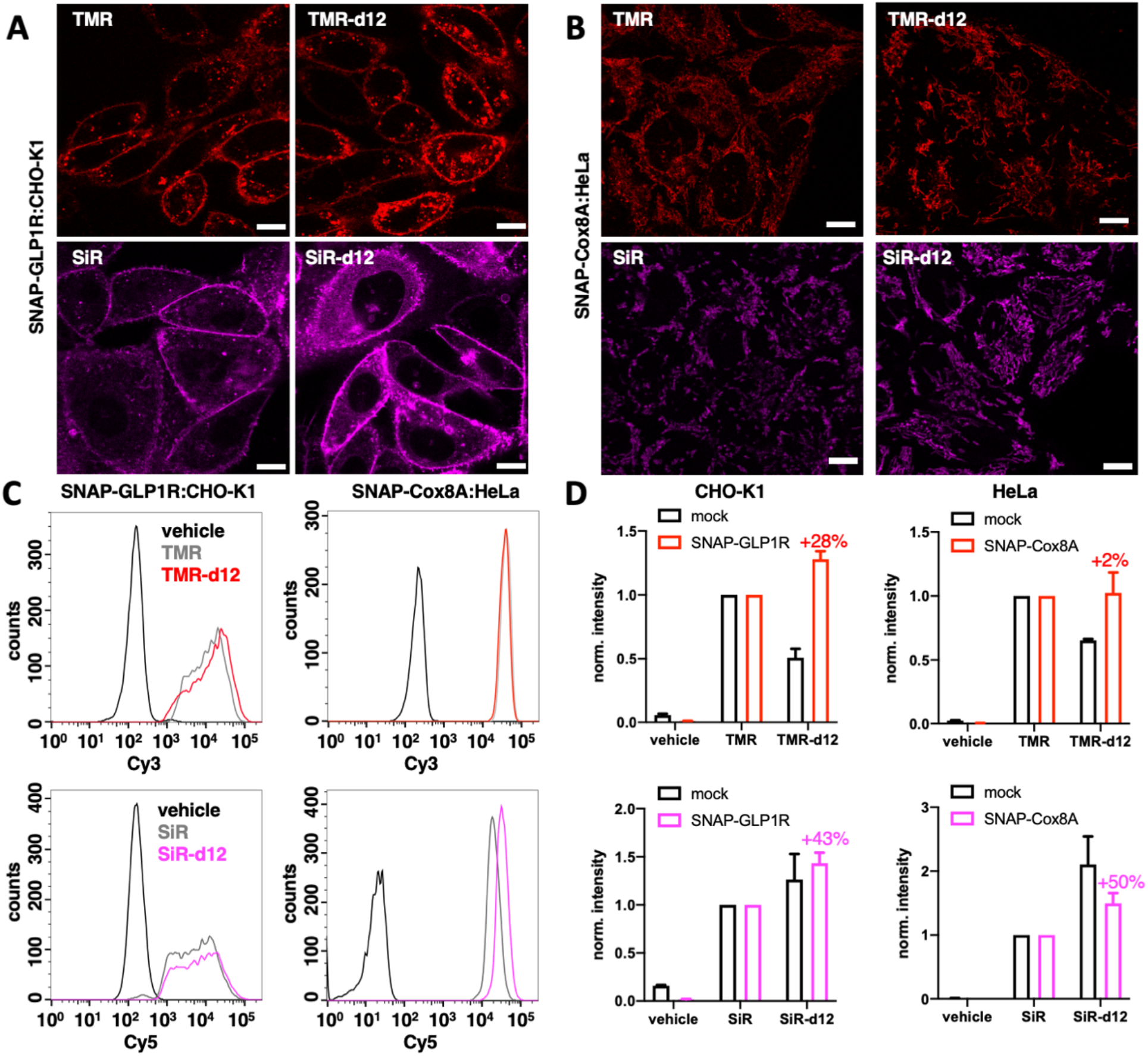
Microscopy and cell sorting reveal increased intensities of rhodamines. A) BG-TMR(-d12) (top) and BG-SiR(-d12) (bottom) label stable SNAP-GLP1R expressing CHO-K1 cells. Scale bar = 10 μm. B) BG-TMR(-d12) (top) and BG-SiR(-d12) (bottom) label stable SNAP-Cox8A expressing HeLa cells. Scale bar = 10 μm. C) Raw data from cell sorting of BG-TMR(-d12) (top) and BG-SiR(-d12) (bottom) labelled stable SNAP-GLP1R:CHO-K1 and SNAP-Cox8A:HeLa cells from A and B. D) As for C, normalized results including mock cells. n=3 independent experiments. Mean±SD.

With these encouraging results, we decided to test our deuterated probes in fluorescent lifetime confocal (FLIM) and stimulated emission by depletion (STED) microscopy, both state-of-the-art imaging techniques to reveal cellular dynamics and structures. Intriguingly, fluorescent lifetimes where higher for d12 congeners than their parent counterparts (τ(TMR) = 2.3 *vs.* τ(TMR-d12) = 2.6 ns; τ(SiR) = 2.9 *vs.* τ(SiR-d12) = 3.5 ns) (Fig. 3A). However, TMR-d12 was not as susceptible to bleaching as TMR, while SiR and SiR-d12 exhibited similar trends of not being prone to fast bleaching in this setup (Fig. 3B). Lastly, and with SiR being one of the most successful far-red dyes for nanoscopy, we investigated super-resolution images acquired in live SNAP-Cox8A:HeLa cells. After incubation with 1 μM BG-SiR or BG-SiR-d12, we recorded images of mitochondrial cristae under the same imaging conditions (Fig. 3C). While both dyes displayed labelling, SiR-d12 was able to resolve cristae sharper with less background, demonstrating its power in nanoscopy.

**Figure 3.**
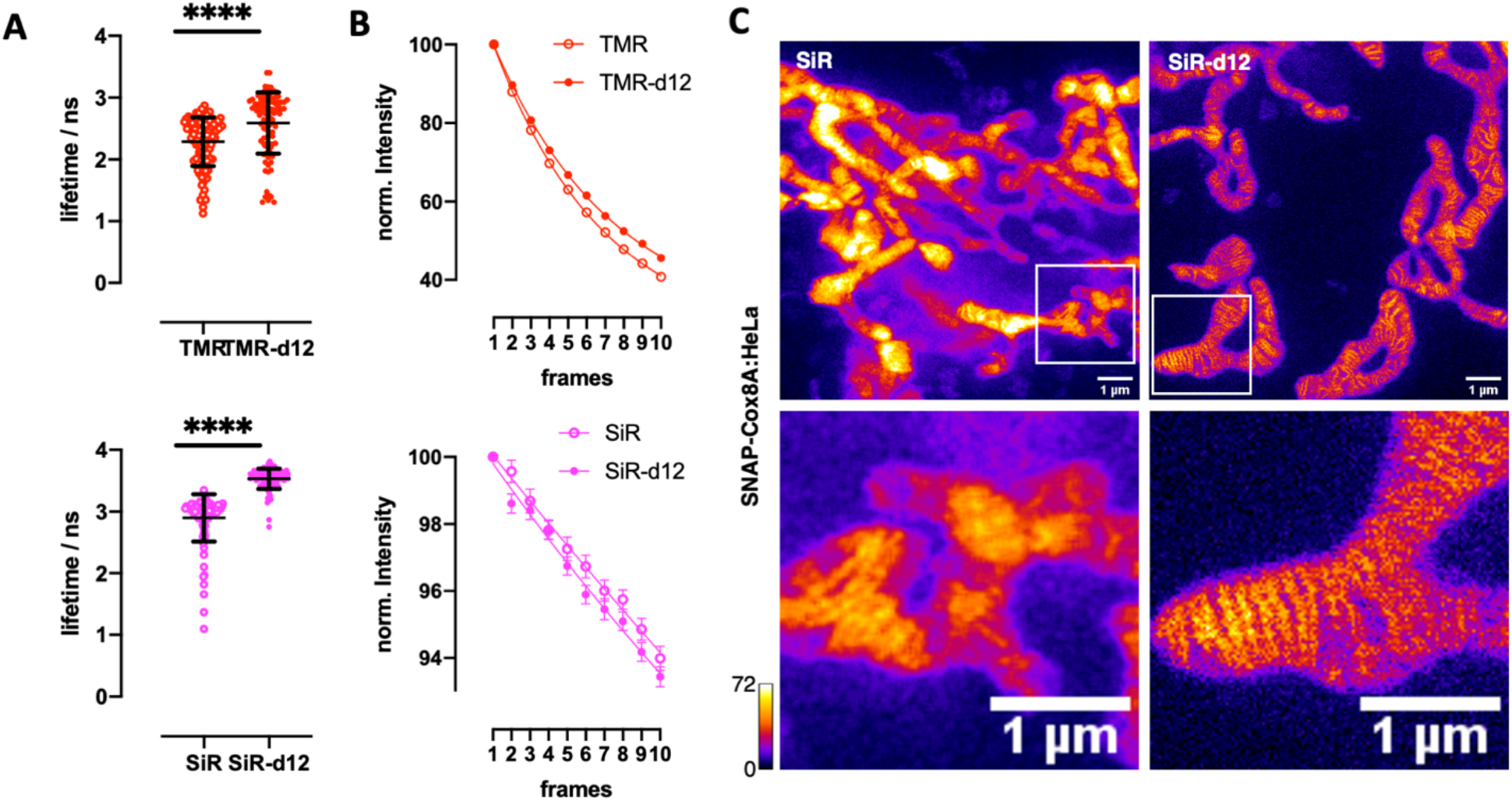
Deuteration increases rhodamine fluorescent lifetimes and shows improved performances and STED nanoscopy. A, B) Fluorescence lifetimes (A) and bleaching curves (B) of BG-TMR(-d12) (top) and BG-SiR(-d12) (bottom) labelled SNAP-GLP1R:CHO-K1 cells. Measurements from individual cells pooled form 2 independent experiments. Mean±SD, **** indicates statistical significance (unpaired t-test, p<0.0001). C) STED nanoscopy of BG-SiR (left) and BG-SiR-d12 (right) in live SNAP-Cox8A:HeLa cells resolving mitochondrial cristae under identical imaging conditions. Scale bar = 1 μm. Representative images of >3 independent experiments.

In our study, we synthesized and tested deuterated fluorophores. We install labelling moieties for SNAP- and Halo-tag conjugation and apply them in different experimental setups, *i.e. in vitro* FRET, and *in cellulo* confocal microscopy, FACS, FLIM and STED. We found that our fluorescent xanthene-based dyes TMR-d12 and SiR-d12 are improved by hydrogen-deuterium exchange on their methyl groups, enhancing photophysical parameters, such as brightness and lifetime, while reducing critical chemical parameters, such as bleaching. While the exact reason for this is unknown, we argue the following: i) affecting the rotation around the aromatic carbon–nitrogen bond (in our case due to higher mass of the CD_3_ groups) has marked effects on fluorescent properties^[17]^, which could suppress non-radiative decays and in turn enhances quantum yield and lifetime^[6]^; and ii) a lower zero-point energy of the C–D *vs.* C–H bond results in slower reaction kinetics, as an higher energy barrier has to be overcome^[18]^, and this could reduce bleaching through for example generated reactive oxygen species. While these explanations need more experimentation, ideally in combination with *in silico* calculations, we showcase novel deuterated dyes that outperform their parent molecules in multiple experiments, ranging from *in vitro* FRET, to live cellular labelling and sorting, in lifetime and super-resolution microscopy. We anticipate this concept i) to be generalizable to more xanthenes (*e.g.* fluoresceins, rhodols, SNARFs, and quenchers like QSYs), and other dye scaffolds (*e.g.* coumarins, cyanines, BODIPYs, EDANS, NBDs, Hoechst) including quenchers at any C–H bond positions for improving and fine-tuning spectroscopic properties; ii) to be further explored with other isotopes, such as ^13^C, ^15^N or even radioactive ^3^H; iii) to be used in different labelling approaches, such as the attachment to sulfonated BG (SBG) scaffolds allowing the separation of SNAP-tagged receptor pools^[19]^, to biomolecule targeting probes^[20–22]^, to “click chemistry” reagents (*e.g.* cyclopropenes, cyclooctenes)^[23]^ or to photoswitchable ligands^[24,25]^ including black hole quenchers (BHQs), and iv) to serve as multimodal dyes for confocal fluorescence and Raman microscopy^[26]^. These efforts are of ongoing interest in our laboratories.

## METHODS

### Synthesis

Chemical synthesis (Supporting Scheme 1) and characterization of compounds is outlined in the Supporting Information. Purity of all dyes was determined to be of >95% by UPLC-UV/Vis traces at 254 nm and dye specific λ_max_ that were recorded on a Waters H-class instrument equipped with a quaternary solvent manager, a Waters autosampler, a Waters TUV detector and a Waters Acquity QDa detector with an Acquity UPLC BEH C18 1.7 μm, 2.1 × 50 mm RP column (Waters Corp., USA).

### Extinction coefficients, Excitation and Emission profiles and quantum yield

To assess photophysical parameters, ^1^H NMR spectra of TMR(d12) and SiR(d12) were recorded with an internal standard (trimethoxybenzene, Aldrich #74588-1g, standard for qNMR) to determine concentrations. The same solutions were diluted 1:1000 in activity buffer (containing in mM: NaCl 50, HEPES 50, pH 7.3) or EtOH + 1% TFA, and absorbance spectra were acquired on a NanodropOne 2000C. Extinction coefficients of d12 dyes were then calculated referenced to literature values of their non-deuterated parental molecules according to equation 1.

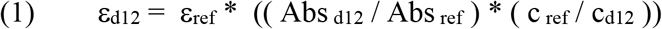

1:100 Dilutions in activity buffer were transferred into Greiner black flat bottom 96 well plates and excitation and emission profiles were recorded on a TECAN INFINITE M PLEX plate reader (TMR: λ_Ex_ = 505±10 nm; λ_Em_ = 550–800±20 nm; 10 flashes; 20 μs integration time; SiR: λ_Ex_ = 605±10 nm; λ_Em_ = 640–800±20 nm; 10 flashes; 20 μs integration time). Absorbance values and integrated emission area (AUC) was used to calculate quantum yield (QY) according to equation 2 under the assumption that there is no change in refractive indices between solutions:

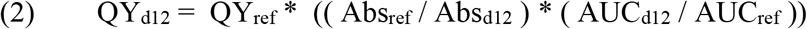

Experiments were run in quadruplicate. Data normalization, integration and plotting was performed in GraphPad Prism 8.

### SNAP_f_ and SNAP–Halo expression and purification

SNAP_f_ was expressed and purified as described previously^[16]^ and complete amino acid sequences for constructs used can be found in the Supporting Information. SNAP-Halo with an N-terminal Strep-tag and C-terminal 10xHis-tag was cloned into a pET51b(+) expression vector for bacterial expression and complete amino acid sequences for constructs used can be found in the Supporting Information. For purification, SNAP-Halo was expressed in the *E. coli* strain BL21 pLysS. LB media contained ampicillin (100 μg/mL) for protein expression. A culture was grown at 37 °C until an OD_600_ of 0.6 was reached at which point cells were induced with IPTG (0.5 mM). Protein constructs were expressed overnight at 16 °C. Cells were harvested by centrifugation and sonicated to produce cell lysates. The lysate was cleared by centrifugation and purified by Ni-NTA resin (Thermofisher) and Strep-Tactin II resin (IBA) according to the manufacturer’s protocols. Purified protein samples were aliquoted in activity buffer (containing in mM: NaCl 50, HEPES 50, pH 7.3), flash frozen and stored at −80 °C.

### SNAP_f_ labelling kinetics

Kinetic measurements were performed on a TECAN Spark Cyto and on a TECAN GENios Pro plate reader by means of fluorescence polarization. Stocks of SNAP_f_ (400 nM) and substrates (100 nM) were prepared in activity buffer (containing in mM: NaCl 50, HEPES 50, pH 7.3) with additional 10 μg/mL BSA. SNAP_f_ and substrates were mixed (100 μL each) in a Greiner black flat bottom 96 well plate. Mixing was performed *via* a built-in injector on a TECAN GENios Pro or by manual pipetting on a TECAN Spark Cyto for TMR and SiR substrates, respectively. Fluorescence polarization reading was started immediately (TMR: λ_Ex_ = 535±25 nm; λ_Em_ = 590±35 nm; 10 flashes; 40 μs integration time; SiR: λ_Ex_ = 605±20 nm; λ_Em_ = 670±20 nm; 10 flashes; 40 μs integration time). Experiments were run in five repetitions and raw polarization values were one-phase decay fitted in GraphPad Prism 8.

### Protein labelling for FRET and full protein mass spectrometry

For protein labelling, 1 μL of the corresponding dye(s) (200 μM in DMSO) were diluted in 220 μL of a 227 nM solution of SNAP-Halo in activity buffer (10 μg/mL BSA was added to controls where no SNAP-Halo protein was present). This resulted in a ∼4-fold excess of labelling substrate and mixing was ensured by carefully pipetting the solution up and down. The reaction mixture was allowed to incubate at r.t. for 1 h before 20 μL were removed for QToF MS analysis to ensure full labelling. The remaining solutions were subjected to spin column purification (Sartorius Vivaspin 500 30 kDa MWCO PES, #VS0122) for three times by adding 500 μL of activity buffer for each cycle. Finally, the solutions were reconstituted in activity buffer, of which 200 μL were transferred into a Greiner black flat bottom 96 well plate and emission spectra were recorded on a TECAN INFINITE M PLEX (TMR: λ_Ex_ = 510±10 nm; λ_Em_ = 550–800±20 nm; 25 flashes; 20 μs integration time; SiR: λ_Ex_ = 610±10 nm; λ_Em_ = 650–800±20 nm; 10 flashes; 20 μs integration time) to observe FRET. Donor or acceptor only labelled constructs, and donor plus acceptor with the addition of 10 μg/mL BSA served as controls, and in these cases acceptor emission was not observed. FRET efficiency was calculated from the sum of maximal emission values derived from the raw spectra I_TMR_ (572–584 nm) and I_SiR_ (660–672 nm) according to equation 3:

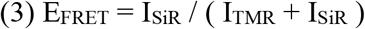

### Cell culture and FACS of SNAP-GLP1R:CHO-K1 and SNAP-Cox8A:HeLa cells

CHO-K1 cells stably expressing the human SNAP-GLP1R (Cisbio) (SNAP-GLP1R:CHO-K1) were maintained in DMEM supplemented with 10% FCS, 1% penicillin/streptomycin, 500 μg/mL G418, 25 mM HEPES, 1% nonessential amino acids and 2% *L*-glutamine. HeLa cells stably expressing SNAP-Cox8A^[15]^ (Cox8A-SNAP:HeLa) were maintained in DMEM supplemented with 10% FCS, 1% penicillin/streptomycin. Cells were incubated with 1 μM of BG-conjugated dye for 30 min at 37 °C, 5% CO_2_, washed with PBS w/o Ca^2+^ and Mg^2+^ and detached using Trypsin/EDTA. Detached cells were pelleted at 300 g at 4 °C, resuspended in DPBS w/o Ca^2+^/Mg^2+^ and kept on ice until fluorescence activated cell sorting (FACS). FACS was performed in LSR-Fortessa (BD Bioscience) on cells gated by SSC/FSC using 561 nm excitation and BP 586/18 (561-586/18 BD channel) for TMR(d12) or 640 nm excitation and BP 670/30 (640-670/30 BD channel) for SiR(d12). Mean fluorescence intensity of >10,000 cells was analyzed and measured using Flow Jo (BD Bioscience).

### FLIM, Confocal and super-resolution microscopy

Live cell fluorescence lifetime and confocal microscopy on SNAP-GLP1R:CHO-K1 was performed using a Leica SP8 with FALCON (Leica Microsystems) equipped with a pulsed white-light excitation laser (80 MHz repetition rate (NKT Photonics)), a 100x objective (HC PL APO CS2 100×/1.40 NA oil), a temperature controlled chamber and operated by LAS X. TMR(d12) were excited using λ = 561 nm and emission signals were captured at λ = 576–670 nm. SiR(d12) were excited using λ = 640 nm and emission signals were captured at λ = 655–748 nm. The emission signals were collected a time gated Hybrid detector (0.5–6 ns). FLIM images of sufficient signal were acquired without gating on Hybrid detectors within 512×512 pxl of 114 nm/pxl with 10 repetitions. Fluorescence lifetime decay curves from selected regions with clear plasma membrane staining were fitted with two exponential functions and the mean amplitude weighted lifetime is reported for each region.

Confocal fluorescence microscopy for photobleaching experiments SNAP-GLP1R were performed on living cells in PBS at room temperature on LSM710 or LSM780 (Carl Zeiss) operated by Zen Black Software using a 63x (1.40 NA oil) objective. TMR(d12) were excited using λ = 561 nm and emission signals were captured at λ =564–712 nm. SiR(d12) were excited using λ = 633 nm and emission signals were captured at λ = 637–740 nm. Emission signal were collected on 34 channel spectral detector (QUASAR, Zeiss), a typical gain of 750 over a scan area of 512×512 pxl.

Confocal and STED microscopy experiments on living Cox8A-SNAP:HelA cells were performed in Life cell imaging buffer (Gibco) at 37 °C using a 100x (1.45 NA lambda oil) objective on a Nikon TiEclipse operated by Micromanager and equipped with a STEDyCON (Abberior Instruments) with 405/488/561/640 excitation and a 775 nm depletion laser. TMR(d12) were excited using λ = 561 nm and emission signals were captured at λ = 580–630 nm. SiR(d12) were excited using λ = 640 nm and emission was collected at λ = 650–700 nm. Both Emission signals were collected by a time gated APD (0.5–8 ns) with 8x signal accumulation and 100 nm pixel size for confocal images. STED images for SiR(d12) were collected with 41% 775nm depletion laser power and 15nm pixel size.

## Acknowledgements

BJ is supported by the Medical Research Council, European Association for the Study of Diabetes, Society for Endocrinology, British Society for Neuroendocrinology and the NIHR Imperial Biomedical Research Centre. We thank Andrea Bergner and Julien Hiblot for providing purified SNAP_f_ protein, Bettina Mathes and Sarah Mikami for synthetic assistance, Cornelia Ulrich and Sebastian Fabritz for mass spectrometry, Kai Johnsson for providing SNAP plasmids, chemical precursors and constant support (all MPIMR). We are grateful to Martina Leidert and Anne Diehl for protein expression and purification, Peter Schmieder for NMR, Ramona Birke and the Hackenberger Laboratory for support (all FMP).

## Author Contributions

JB conceived and supervised the study. JB designed, synthesized and characterized chemical compounds. KR, CC and JB performed *in vitro* measurements. KCA, JE and ML performed cell culture, cell sorting and microscopy. BJ provided reagents. JB wrote the manuscript with input from all authors.

