## Supporting Information for "Deuterated rhodamines for protein labelling in nanoscopy"

**Table of Contents**

**1. General 3**

**2. Synthesis 5**

*2.1. General procedure A for fluorophore coupling 5*

*2.2. 3-(Bis(methyl-$d_3$)amino)phenol (2) 5*

*2.3. 2-(6-(Bis(methyl-$d_3$)amino)-3-(bis(methyl-$d_3$)iminio)-3H-xanthen-9-yl)-4-carboxybenzoate (3) 6*

*2.4. 4-((4-(((2-Amino-9H-purin-6-yl)oxy)methyl)benzyl)carbamoyl)-2-(6-(bis(methyl-$d_3$)amino)-3-(bis(methyl-$d_3$)iminio)-3H-xanthen-9-yl)benzoate (BG-TMR-d12) 7*

*2.5. 2-(6-(Bis(methyl-$d_3$)amino)-3-(bis(methyl-$d_3$)iminio)-3H-xanthen-9-yl)-4-((2-(2-((6-chlorohexyl)oxy)ethoxy)ethyl)carbamoyl)benzoate (Halo-TMR-d12) 8*

*2.6. 3,7-Diamino-5,5-dimethyl-3'-oxo-3'H,5H-spiro[dibenzo[b,e]siline-10,1'-isobenzofuran]-6'-carboxylic acid (5) 9*

*2.7. 3,7-Bis(bis(methyl-$d_3$)amino)-5,5-dimethyl-3'-oxo-3'H,5H-spiro[dibenzo[b,e]siline-10,1'-isobenzofuran]-6'-carboxylic acid (6) 10*

*2.8. 4-((4-(((2-Amino-9H-purin-6-yl)oxy)methyl)benzyl)carbamoyl)-2-(7-(bis(methyl-$d_3$)amino)-3-(bis(methyl-$d_3$)iminio)-5,5-dimethyl-3,5-dihydrodibenzo[b,e]silin-10-yl)benzoate (BG-SiR-d12) 11*

*2.9. 2-(7-(Bis(methyl-$d_3$)amino)-3-(bis(methyl-$d_3$)iminio)-5,5-dimethyl-3,5-dihydrodibenzo[b,e]silin-10-yl)-4-((2-(2-((6-chlorohexyl)oxy)ethoxy)ethyl)carbamoyl)benzoate (Halo-SiR-d12) 12*

**3. $SNAP_f$ construct 13**

**4. SNAP–Halo construct and mass spectrometry 13**

**5. Supplementary Schemes 14**

**6. References 15**

### 1. General

All chemical reagents and anhydrous solvents for synthesis were purchased from commercial suppliers (Sigma-Aldrich, Fluka, Acros, Fluorochem, TCI) and were used without further purification. BG-TMR and BG-SiR were described before.[1]

NMR spectra were recorded in deuterated solvents on a Bruker AVANCE III 600 equipped with a CryoProbe or on a Bruker AVANCE II 750 and calibrated to residual solvent peaks ($^{1}H/^{13}C$ in ppm): DMSO-$d_6$ (2.50/39.52), MeOD-$d_4$ (3.31/49.00). Multiplicities are abbreviated as follows: s = singlet, d = doublet, t = triplet, q = quartet, p = pentet, h = heptet, br = broad, m = multiplet. Coupling constants *J* are reported in Hz. Spectra are reported based on appearance, not on theoretical multiplicities derived from structural information. DCl was added to some TMR compounds to improve line sharpening and resolution: 2 µL of 35% DCl was diluted in 198 µL $D_2O$ to obtain a 0.35% solution, of which 30 µL were added to 570 µL of DMSO-$d_6$.

UPLC-UV/Vis for purity assessment was performed on a Waters H-class instrument equipped with a quaternary solvent manager, a Waters autosampler, a Waters TUV detector and a Waters Acquity QDa detector with an Acquity UPLC BEH C18 1.7 µm, 2.1 × 50 mm RP column (Waters Corp., USA). Buffer A: 0.1% TFA in $H_2O$ Buffer B: 0.1% TFA in MeCN. The typical gradient was from 5% B for 0.5 min; gradient to 95% B over 3.0 min; 95% B for 0.9 min; gradient to 5% B over 1.1 min with 0.6 mL/min flow. Chromatograms were loaded into Graphpad Prism8 and purity was determined by calculating AUC ratios.

High resolution mass spectrometry was performed using a Bruker maXis II ETD hyphenated with a Shimadzu Nexera system. The instruments were controlled via Brukers otofControl 4.1 and Hystar 4.1 SR2 (4.1.31.1) software. The acquisition rate was set to 3 Hz and the following source parameters were used for positive mode electrospray ionization: End plate offset = 500 V; capillary voltage = 3800 V; nebulizer gas pressure = 45 psi; dry gas flow = 10 L/min; dry temperature = 250 °C. Transfer, quadrupole and collision cell settings are mass range dependent and were fine-adjusted with consideration of the respective analyte's molecular weight. For internal calibration sodium format clusters were used. Samples were desalted via fast liquid chromatography. A Supelco TitanTM C18 UHPLC Column, 1.9 µm, 80 Å pore size, 20 × 2.1 mm and a 2 min gradient from 10 to 98% aqueous MeCN with 0.1% FA ($H_2O$: Carl Roth GmbH + Co. KG ROTISOLV® Ultra LC-MS; MeCN: Merck KGaA LiChrosolv® Acetonitrile hypergrade for LC-MS; FA - Merck KGaA LiChropur® Formic acid 98%–100% for LC-MS) was used for separation. Sample dilution in 10% aqueous MeCN (hyper grade) and injection volumes were chosen dependent of the analyte's ionization efficiency. Hence, on-column loadings resulted between 0.25–5.0 ng. Automated internal re-calibration and data analysis of the recorded spectra were performed with Bruker's DataAnalysis 4.4 SR1 software.

Intact proteins were analyzed using a Waters H-class instrument equipped with a quaternary Solvent manager, a Waters sample manager-FTN, a Waters PDA detector and a Waters column manager with an Acquity UPLC protein BEH C4 column (300 Å, 1.7 µm, 2.1 mm x 50 mm). Proteins were eluted with a flow rate of 0.3 mL/min at a column temperature of 80 °C. The following gradient was used: A: 0.01% FA in $H_2O$; B: 0.01% FA in MeCN. gradient 5-95% B from 0-6 min. Mass analysis was conducted with a Waters XEVO G2-XS QTof analyzer. Proteins were ionized in positive ion mode applying a cone voltage of 40 kV. Raw data was analyzed with MaxEnt 1. After deconvolution of the crude spectra, no single or non-labelled SNAP-Halo construct was observed, indicating complete reaction.

Preparative RP-HPLC was performed on a Waters e2695 system equipped with a 2998 PDA detector for product collection (550, 600 or 650 nm) on a Supelco Ascentis® C18 HPLC Column (5 µm, 250 × 21.2 mm). Buffer A: 0.1% TFA in $H_2O$ Buffer B: MeCN. The typical gradient was from 10% B for 5 min; gradient to 90% B over 45 min to 90% B for 5 min; gradient to 99% B over 5 min with 8 mL/min flow.

Flash column chromatography (FCC) was performed on a Biotage Isolera One with pre-packed silica columns (0.040–0.063 mm, 230-400 mesh, Silicycle). Reactions and chromatography fractions were monitored by thin layer chromatography (TLC) on Merck silica gel 60 F254 glass plates. The spots were visualized either under UV light at 254 nm and/or 366 nm or with appropriate staining method (iodine, *para*-anisaldehyde, $KMnO_4$) followed by heating.

### 2. Synthesis

#### 2.1. General procedure A for fluorophore coupling

In an Eppendorf tube, 1.0 equiv. of deuterated carboxy-dye was dissolved in 100 µL/mg DMSO and 8.0 equiv. of DIPEA. Upon addition of 1.5 equiv. TSTU (from a 10 mg/mL stock in DMSO) the reaction mixture was vortexed and allowed to incubate for 10 min, before 1.5 equiv. of amine (BG-$NH_2$ or Halo-$NH_2$) were added. The mixture was vortexed again and allowed to incubate for 60 min before it was quenched by addition of 20 equiv. of acetic acid and 25vol% of water. HPLC (MeCN:$H_2O$+0.1% TFA = 10:90 to 90:10 over 60 minutes) provided the desired compound, which was aliquoted to 5 nmol and obtained as a colorful powders after lyophilization.

#### 2.2. 3-(Bis(methyl-d$_3$)amino)phenol (2)

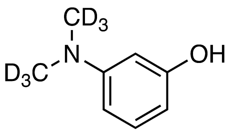


A round bottom flask was charged with 1.00 g (9.00 mmol, 1.0 equiv.) of 3-aminophenol (**1**) and dissolved in 50 mL of EtOH before 2.49 g (18.0 mmol, 2.0 equiv.) of $K_2CO_3$ and 2.67 g (18.4 mmol, 1.15 mL, 2.05 equiv.) iodomethane-d$_3$ were added. The reaction mixture was heated to 80 °C o.n. before it was cooled to r.t. and all volatiles were removed *in vacuo*. FCC (10% EtOAc in hexanes) provided 763 mg (5.33 mmol) of the desired compound as a beige powder in 59% yield.

**$^1$H NMR** (400 MHz, $CDCl_3$): $\delta$ [ppm] = 7.09 (t, $J$ = 8.1 Hz, 1H), 6.36–6.29 (m, 1H), 6.27–6.18 (m, 2H).
**$^{13}$C NMR** (100 MHz, $CDCl_3$): $\delta$ [ppm] = 156.6, 152.0, 130.0, 105.5, 103.9, 99.8, 39.8 (m$_c$, $J$ = 20 Hz).
**HRMS** (ESI): calc. for $C_8H_6D_6NO$ $[M+H]^+$: 144.1290, found: 144.1290.

#### 2.3. 2-(6-(Bis(methyl-$d_3$)amino)-3-(bis(methyl-$d_3$)iminio)-3*H*-xanthen-9-yl)-4-carboxybenzoate (3)

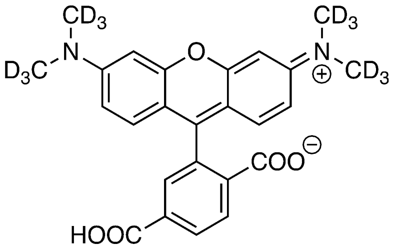


A round bottom flask was charged with 200 mg (1.40 mmol, 1.0 equiv.) of **2** and dissolved in 5 mL of toluene before 322 mg (1.68 mmol, 1.2 equiv.) of 1,2,4-benzenetricarboxylic anhydride was added. The reaction mixture was heated to 130 °C o.n. before it was cooled to r.t., MeOH and celite were added and all volatiles were removed *in vacuo*. Dry load FCC (DCM:MeOH = 100:0 to 50:50 over 45 CV) provided crude TMR-d12. HPLC (MeCN:$H_2O$+0.1% TFA = 30:70 to 70:30 over 60 minutes, first red fraction) was performed to separate isomers and to provided 19.8 mg (44.8 µmol) of the desired compound as a red powder in 6% yield.

Among this, we also isolated: 4-(4-(bis(methyl-$d_3$)amino)-2-hydroxybenzoyl)isophthalic acid, 2-(4-(bis(methyl-$d_3$)amino)-2-hydroxybenzoyl)terephthalic acid, 2-(6-(bis(methyl-$d_3$)amino)-3-(bis(methyl-$d_3$)iminio)-3H-xanthen-9-yl)-5-carboxybenzoate.

**$^1$H NMR** (600 MHz, MeOD-$d_4$): $\delta$ [ppm] = 8.40 (d, $J$ = 8.1 Hz, 1H), 8.38 (dd, $J$ = 8.2, 1.5 Hz, 1H), 7.16 (d, $J$ = 9.5 Hz, 2H), 7.06 (dd, $J$ = 9.5, 2.5 Hz, 3H), 6.98 (d, $J$ = 2.4 Hz, 2H).
**$^{13}$C NMR** (150 MHz, MeOD-$d_4$): $\delta$ [ppm] = 167.9, 167.7, 160.6, 159.2, 159.1, 136.7, 135.9, 135.3, 132.7, 132.3, 132.2, 132.0, 115.5, 114.9, 97.4, 40.1 (h, $J$ = 21.1 Hz).
**HRMS** (ESI): calc. for $C_{25}H_{11}D_{12}N_2O_5$ $[M+H]^+$: 443.2355, found: 443.2352.

### 2.4. 4-((4-(((2-Amino-9*H*-purin-6-yl)oxy)methyl)benzyl)carbamoyl)-2-(6-(bis(methyl-$d_3$)amino)-3-(bis(methyl-$d_3$)iminio)-3*H*-xanthen-9-yl)benzoate (BG-TMR-d12)

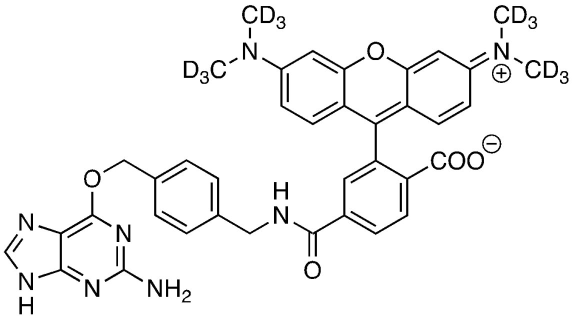

**BG–TMR–d12** was prepared according to general procedure A.

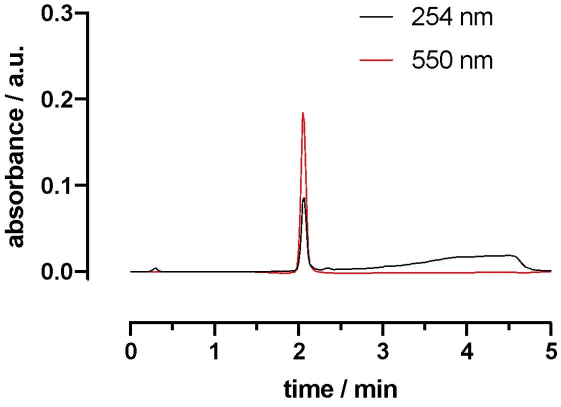


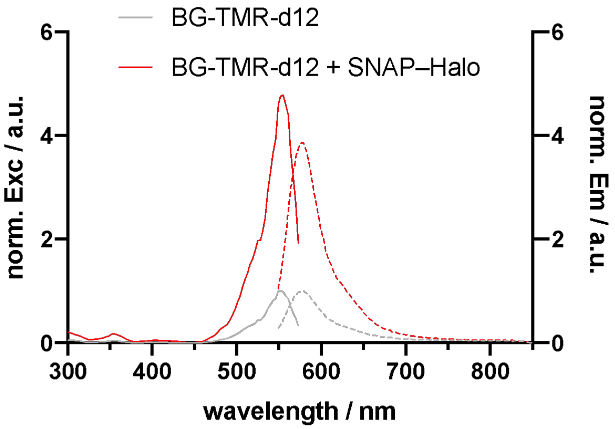


**$^1$H NMR** (600 MHz, DCl in DMSO-$d_6$): $\delta$ [ppm] = 8.62 (s, 1H), 8.31 (d, $J$ = 8.2 Hz, 1H), 8.26 (dd, $J$ = 8.3, 1.9 Hz, 1H), 7.89 (d, $J$ = 1.9 Hz, 1H), 7.50 (d, $J$ = 7.9 Hz, 2H), 7.37 (d, $J$ = 7.9 Hz, 2H), 7.06 (dd, $J$ = 9.5, 2.4 Hz, 2H), 7.03 (d, $J$ = 9.5 Hz, 2H), 6.96 (d, $J$ = 2.4 Hz, 2H), 5.54 (s, 2H), 4.48 (s, 2H).
**HRMS** (ESI): calc. for $C_{38}H_{24}D_{12}N_8O_5$ $[M+2H]^{2+}$: 348.1776, found: 348.1770.

### 2.5. 2-(6-(Bis(methyl-d$_3$)amino)-3-(bis(methyl-d$_3$)iminio)-3*H*-xanthen-9-yl)-4-((2-(2-((6-chlorohexyl)oxy)ethoxy)ethyl)carbamoyl)benzoate (Halo-TMR-d12)

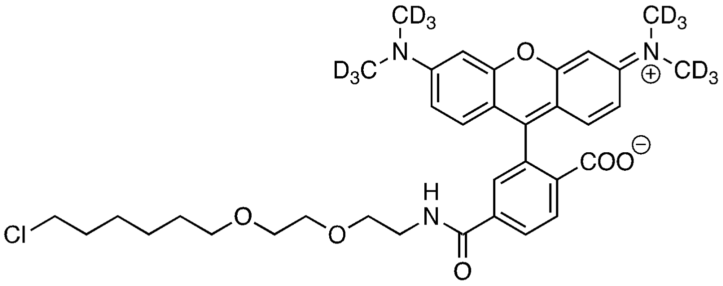


**Halo–TMR–d12** was prepared according to general procedure A.

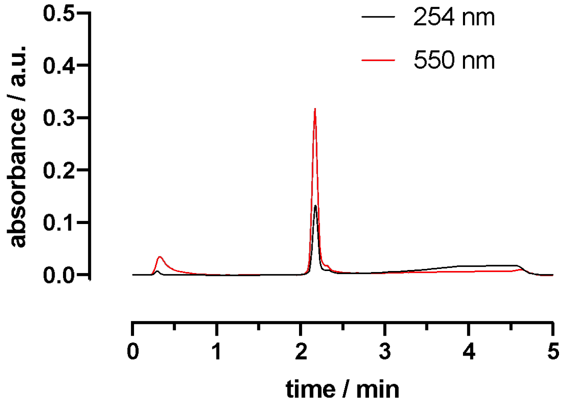


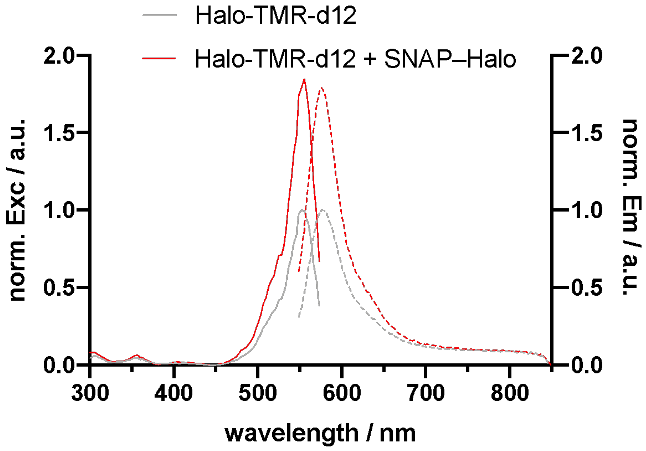


**$^1$H NMR** (600 MHz, DCl in DMSO-d$_6$): $\delta$ [ppm] = 8.29 (d, $J$ = 8.1 Hz, 1H), 8.22 (dt, $J$ = 8.3, 1.8 Hz, 1H), 7.85–7.79 (m, 1H), 7.07 (dd, $J$ = 9.5, 2.4 Hz, 2H), 7.03 (dd, $J$ = 9.6, 2.0 Hz, 2H), 6.96 (d, $J$ = 2.3 Hz, 2H), 3.42 (dt, $J$ = 11.9, 5.5 Hz, 5H), 3.31 (t, $J$ = 6.5 Hz, 2H), 1.65 (p, $J$ = 6.7 Hz, 2H), 1.40 (p, $J$ = 6.7 Hz, 2H), 1.36–1.27 (m, 2H), 1.28–1.12 (m, 4H). 4 protons presumably masked by $D_2O$.

**HRMS** (ESI): calc. for $C_{35}H_{31}D_{12}ClN_3O_6$ $[M+H]^+$: 648.3588, found: 648.3587.

### 2.6. 3,7-Diamino-5,5-dimethyl-3'-oxo-3'*H*,5*H*-spiro[dibenzo[*b,e*]siline-10,1'-isobenzofuran]-6'-carboxylic acid (5)

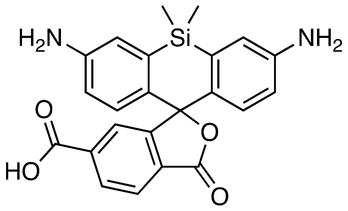


A Schlenk flask was charged under an argon atmosphere with 50.0 mg (67.7 µmol, 1.0 equiv.) of *tert*-butyl 5,5-dimethyl-3'-oxo-3,7-bis(((trifluoromethyl)sulfonyl)oxy)-3'*H*,5*H*-spiro[dibenzo[*b,e*]-siline-10,1'-isobenzo-furan]-6'-carboxylate[3] (**4**), 12.4 mg (13.5 µmol, 0.2 equiv.) of tris(dibenzylideneacetone)dipalladium(0) ($Pd_2dba_3$), 9.7 mg (20.3 µmol, 0.3 equiv.) of 2-(dicyclohexylphosphino)-2',4',6'-tri*iso*propylbiphenyl (XPhos), 106 mg (325 µmol, 4.8 equiv.) of $Cs_2CO_3$ and 36.8 mg (203 µmol, 34.1 µL, 3.0 equiv.) of benzophenone imine ($Ph_2CNH$), before 2 mL of dry 1,4-dioxane were added via syringe. The reaction mixture was heated to 100 °C o.n. before it was cooled to r.t. and all volatiles were removed *in vacuo*. 5% TFA in $dH_2O$ was added to the crude and the deprotection step was allowed to incubate at r.t. over 4 hours. HPLC (MeCN:$H_2O$+0.1% TFA = 10:90 to 90:10 over 60 minutes) provided 22.2 mg (53.4 µmol) of the desired compound as a purple-blue powder in 79% yield.

**$^1$H NMR** (400 MHz, MeOD-$d_4$): $\delta$ [ppm] = 8.26 (dd, $J$ = 8.1, 1.4 Hz, 1H), 8.11 (d, $J$ = 8.1 Hz, 1H), 7.85 (dd, $J$ = 1.5, 0.7 Hz, 1H), 7.26 (d, $J$ = 2.5 Hz, 2H), 6.86 (d, $J$ = 8.8 Hz, 2H), 6.75 (dd, $J$ = 8.8, 2.4 Hz, 2H), 0.65 (s, 3H), 0.56 (s, 3H).
**HRMS** (ESI): calc. for $C_{23}H_{21}N_2O_4Si$ $[M+H]^+$: 417.1265, found: 417.1260.

#### 2.7. 3,7-Bis(bis(methyl-d$_3$)amino)-5,5-dimethyl-3'-oxo-3'*H*,5*H*-spiro[dibenzo[*b,e*]siline-10,1'-isobenzofuran]-6'-carboxylic acid (6)

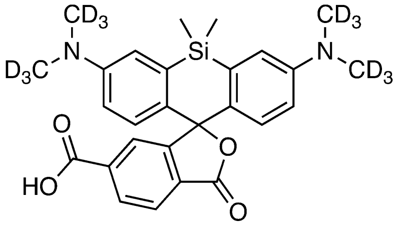

A round bottom flask was charged with 20.5 mg (49.2 µmol, 1.0 equiv.) of **5** dissolved in 2 mL of EtOH before 13.6 mg (98.4 µmol, 2.0 equiv.) of $K_2CO_3$ and 57.8 mg (394 µmol, 24.5 µL, 8.0 equiv.) iodomethane-d$_3$ were added. The reaction mixture was heated to 80 °C o.n. before it was cooled to r.t. and all volatiles were removed *in vacuo*. HPLC (MeCN:$H_2O$+0.1% TFA = 10:90 to 90:10 over 60 minutes) provided 5.2 mg (10.7 µmol) of the desired compound as a blue powder in 22% yield.

**$^1$H NMR** (600 MHz, DMSO-d$_6$): $\delta$ [ppm] = 8.11 (dd, J = 8.0, 1.3 Hz, 1H), 8.04 (d, J = 8.0, 1H), 7.63 (d, J = 1.3 Hz, 1H), 7.01 (d, J = 2.8, 2H), 6.69 (d, J = 8.9 Hz, 2H), 6.65 (dd, J = 9.0, 2.9 Hz, 2H), 0.64 (s, 3H), 0.54 (s, 3H).
**HRMS** (ESI): calc. for $C_{27}H_{17}D_{12}N_2O_4Si$ $[M+H]^+$: 485.2644, found: 485.2645.

#### 2.8. 4-((4-(((2-Amino-9*H*-purin-6-yl)oxy)methyl)benzyl)carbamoyl)-2-(7-(bis(methyl-d$_3$)amino)-3-(bis(methyl-d$_3$)iminio)-5,5-dimethyl-3,5-dihydrodibenzo[*b,e*]silin-10-yl)benzoate (BG-SiR-d12)

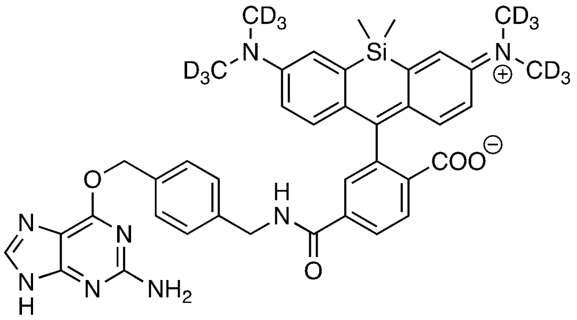


**BG–SiR–d12** was prepared according to general procedure A.

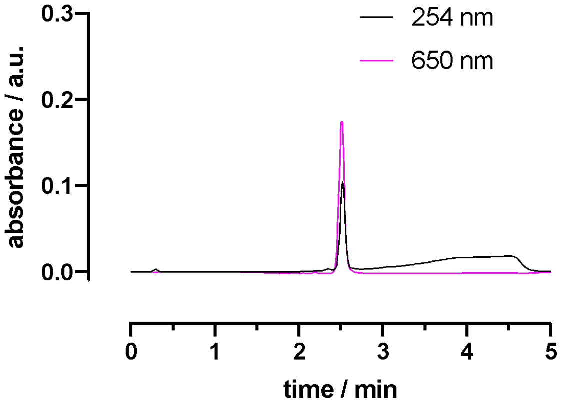


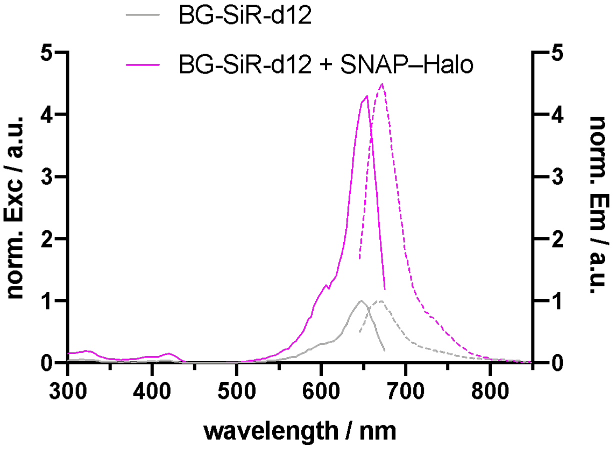


**$^1$H NMR** (600 MHz, DMSO-d$_6$): $\delta$ [ppm] = 9.29 (t, $J$ = 6.0 Hz, 1H), 8.12 (dd, $J$ = 8.1, 1.4 Hz, 1H), 8.03 (d, $J$ = 8.1 Hz, 1H), 7.70 (d, $J$ = 1.3 Hz, 1H), 7.44 (d, $J$ = 7.9 Hz, 2H), 7.31 (d, $J$ = 8.0 Hz, 2H), 7.00 (d, $J$ = 2.5 Hz, 2H), 6.64 (dd, $J$ = 9.0, 2.6 Hz, 2H), 6.62 (d, $J$ = 9.6 Hz, 2H), 6.27 (s, 2H), 5.44 (s, 2H), 4.44 (d, $J$ = 5.8 Hz, 2H), 0.62 (s, 3H), 0.51 (s, 3H).
**HRMS** (ESI): calc. for $C_{40}H_{30}D_{12}N_8O_4Si$ [M+2H]$^{2+}$: 369.1920, found: 369.1917.

#### 2.9. 2-(7-(Bis(methyl-d3)amino)-3-(bis(methyl-d3)iminio)-5,5-dimethyl-3,5-dihydrodibenzo[*b,e*]silin-10-yl)-4-((2-(2-((6-chlorohexyl)oxy)ethoxy)ethyl)carbamoyl)benzoate (Halo-SiR-d12)

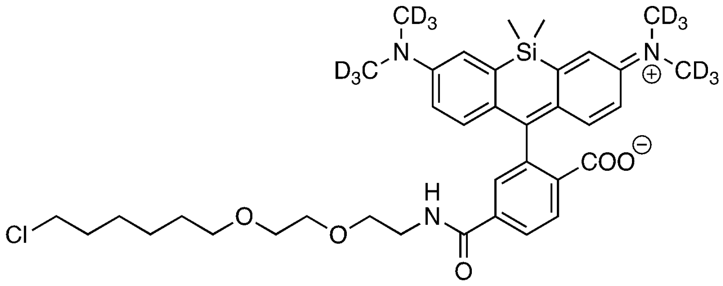

**Halo–SiR–d12** was prepared according to general procedure A.

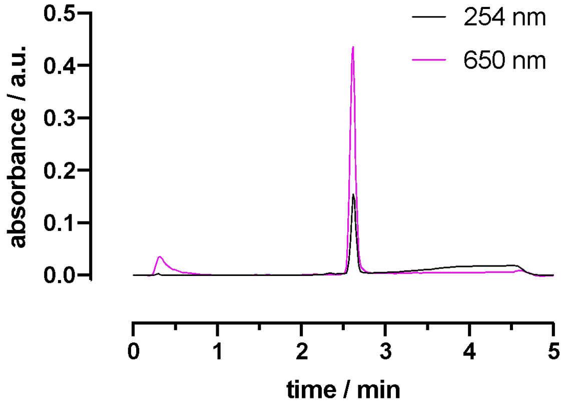


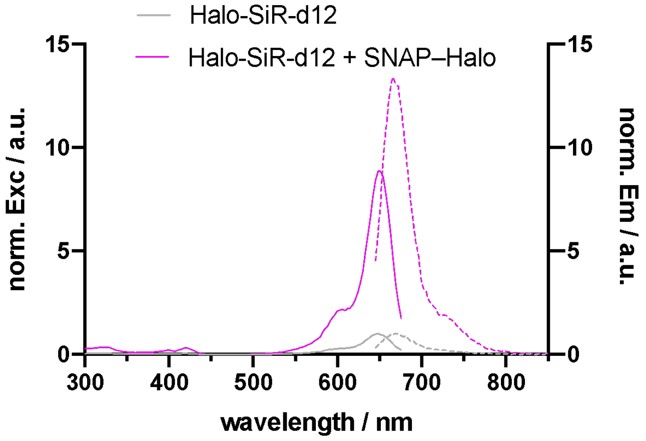


**[1]H NMR** (600 MHz, DMSO-d6): $\delta$ [ppm] = 8.77 (t, $J$ = 5.6 Hz, 1H), 8.08 (dd, $J$ = 8.1, 1.4 Hz, 1H), 8.02 (d, $J$ = 8.0 Hz, 1H), 7.67 (d, $J$ = 1.3 Hz, 1H), 7.01 (d, $J$ = 2.7 Hz, 2H), 6.63 (dd, $J$ = 8.9, 2.7 Hz, 2H), 6.61 (d, $J$ = 8.9 Hz, 2H), 3.57 (t, $J$ = 6.6 Hz, 2H), 3.49 (dt, $J$ = 5.4, 1.9 Hz, 4H), 3.43 (dd, $J$ = 5.9, 3.7 Hz, 2H), 3.37 (q, $J$ = 5.9 Hz, 2H), 2.54–2.51 (m, 4H), 1.65 (dt, $J$ = 14.3, 6.8 Hz, 2H), 1.41 (dt, $J$ = 14.1, 6.7 Hz, 2H), 1.34–1.28 (m, 2H), 0.64 (s, 3H), 0.52 (s, 3H).
**HRMS** (ESI): calc. for $C_{37}H_{37}D_{12}ClN_3O_5Si$ $[M+H]^+$: 690.3878, found: 690.3874.

### 3. SNAP$_f$ construct

SNAP$_f$ sequence:

```
MASWSHPQFE KGADDDDKVP HMDKDCEMKR TTLDSPLGKL ELSGCEQGLH RIIFLGKGTS
AADAVEVPAP AAVLGGPEPL MQATAWLNAY FHQPEAIEEF PVPALHHPVF QQESFTRQVL
WKLLKVVKFG EVISYSHLAA LAGNPAATAA VKTALSGNPV PILIPCHRVV QGDLDVGGYE
GGLAVKEWLL AHEGHRLGKP GLGAPGFSSI SAHHHHHHHHHH
```

Strep-Tag II, Enterokinase-site, SNAP$_f$, His-Tag

### 4. SNAP–Halo construct and mass spectrometry

SNAP–Halo sequence:

```
MASWSHPQFE KGADDDDKVP HMDKDCEMKR TTLDSPLGKL ELSGCEQGLH EIIFLGKGTS
AADAVEVPAP AAVLGGPEPL MQATAWLNAY FHQPEAIEEF PVPALHHPVF QQESFTRQVL
WKLLKVVKFG EVISYSHLAA LAGNPAATAA VKTALSGNPV PILIPCHRVV QGDLDVGGYE
GGLAVKEWLL AHEGHRLGKP GLGGRLEVLF QGPKAFLEGS EIGTGFPFDP HYVEVLGERM
HYVDVGPRDG TPVLFLHGNP TSSYVWRNII PHVAPTHRCI APDLIGMGKS DKPDLGYFFD
DHVRFMDAFI EALGLEEVVL VIHDWGSALG FHWAKRNPER VKGIAFMEFI RPIPTWDEWP
EFARETFQAF RTTDVGRKLI IDQNVFIEGT LPMGVVRPLT EVEMDHYREP FLNPVDREPL
WRFPNELPIA GEPANIVALV EEYMDWLHQS PVPKLLFWGT PGVLIPPAEA ARLAKSLPNC
KAVDIGPGLN LLQEDNPDLI GSEIARWLST LEISGAPGFS SISAHHHHHH HHHH*
```

Strep-Tag II, Enterokinase-site, SNAP, Precission Sequence, Halo, His-Tag

| Condition | calc. | found |
|---|---|---|
| SNAP–Halo | 59072 | 59069 |
| SNAP–Halo:BG-TMR+Halo-SiR | 60241 | 60245 |
| SNAP–Halo:BG-SiR+Halo-TMR | 60241 | 60241 |
| SNAP–Halo:BG-TMR-d12+Halo-SiR-d12 | 60265 | 60269 |
| SNAP–Halo:BG-SiR-d12+Halo-TMR-d12 | 60265 | 60267 |
| SNAP–Halo:BG-TMR | 59600 | 59602 |
| SNAP–Halo:BG-SiR | 59645 | 59642 |
| SNAP–Halo:Halo-TMR | 59671 | 59669 |
| SNAP–Halo:Halo-SiR | 59713 | 59711 |
| SNAP–Halo:BG-TMR-d12 | 59612 | 59615 |
| SNAP–Halo:BG-SiR-d12 | 59657 | 59656 |
| SNAP–Halo:Halo-TMR-d12 | 59683 | 59682 |
| SNAP–Halo:Halo-SiR-d12 | 59725 | 59724 |

### 5. Supplementary Schemes

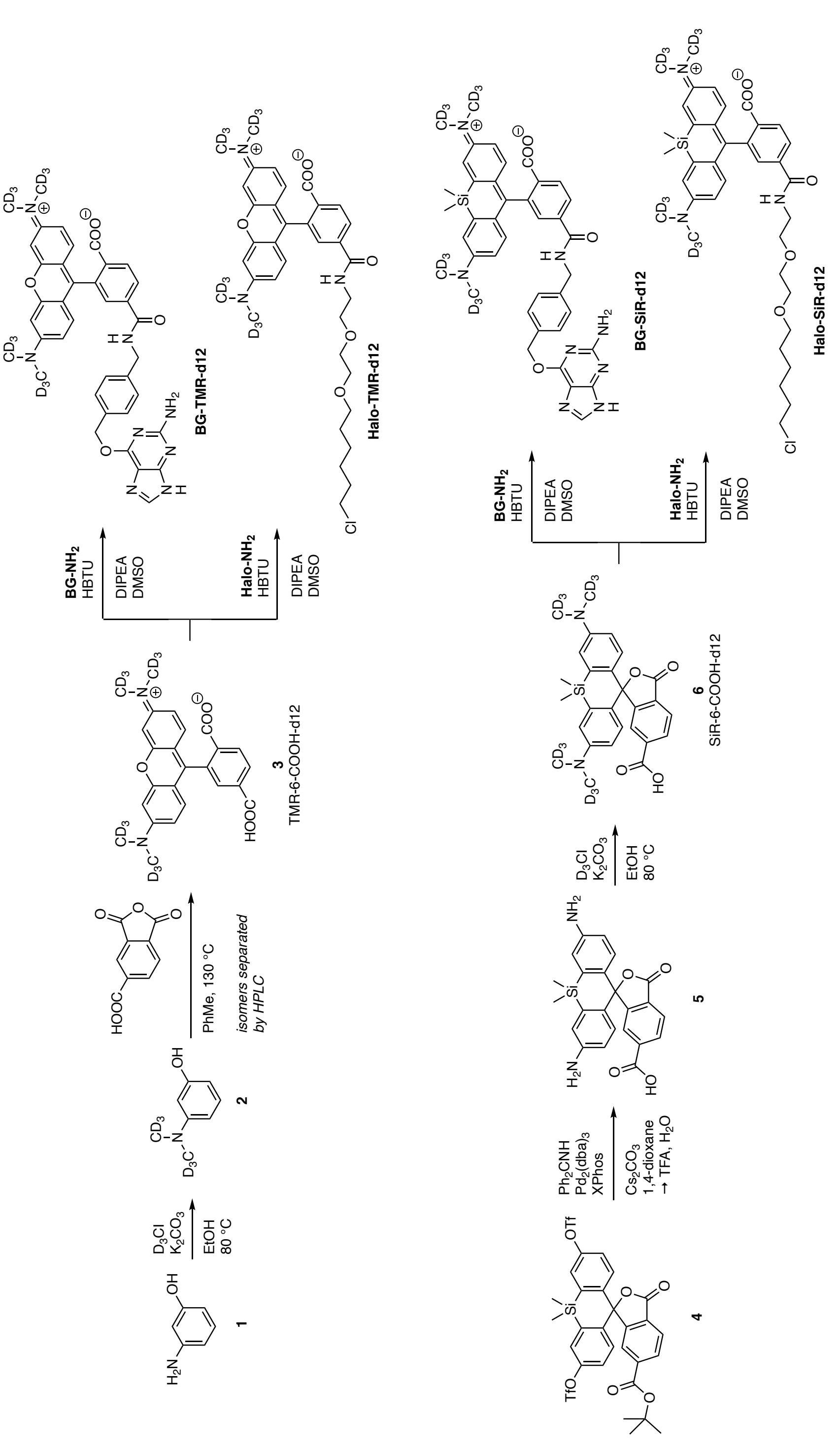


**Supporting Scheme 1: Chemical synthesis of d12 probes.**
